# Functional connectivity of thalamic nuclei during sensorimotor task-based fMRI at 9.4 Tesla

**DOI:** 10.1101/2025.02.09.632791

**Authors:** Edyta Charyasz, Michael Erb, Jonas Bause, Rahel Heule, Benjamin Bender, Vinod Jungir Kumar, Wolfgang Grodd, Klaus Scheffler

## Abstract

The thalamus is the brain’s central communication hub, playing a key role in processing and relaying sensorimotor and cognitive information between the cerebral cortex and other brain regions. It consists of specific and non-specific nuclei, each with a different role. Specific thalamic nuclei relay sensory and motor information to specific cortical and subcortical regions to ensure precise communication. In contrast, non-specific thalamic nuclei are involved in general functions such as attention or consciousness through broader and less targeted connections. In the present study, we aimed to investigate the functional connectivity patterns of the thalamic nuclei identified in our previous study as being involved in motor (finger-tapping) and sensory (finger-touch) tasks. The results of this study show that thalamic nuclei are not static hubs with a predefined role in neural signal processing, as they show different task-specific functional connectivity patterns in the anterior, middle, lateral, and posterior thalamic nuclei. Instead, they are all functional hubs that can flexibly change their connections to other brain regions in response to task demands. This work has important implications for understanding task-dependent functional connectivity between thalamic nuclei and different brain regions using task-based fMRI at 9.4 Tesla.

## 1. INTRODUCTION

The human brain is a complex and interconnected network of approximately 86 billion neurons (Azevedo et al., 2009), where coordinated activity between different brain regions enables functional processing in the brain. The cerebral cortex, for instance, is responsible for sensory processing, motor coordination, balance, motor learning and higher-level cognitive tasks such as memory, language, perception, and emotional regulation (Rolls, 2016). Similarly, the cerebellum is involved in motor coordination, balance, motor learning, cognitive processes, and emotional regulation (Manto et al., 2012, De Zeeuw and Ten Brinke, 2015, Prati et al., 2024, Rudolph et al., 2023). However, a key role in subcortical-cortical and cortico-cortical functional communication is played by the thalamus, a small structure located deep in the brain that relays sensory, cognitive and motor signals throughout the brain (Sherman and Guillery, 2006, Shine et al., 2023, Haber and McFarland, 2001, Sherman, 2007). The thalamus is composed of several distinct nuclei with specific and non-specific connections to cerebral and cerebellar areas (Ward, 2013, Jones, 1983, Zhou et al., 2011, Jones, 1998). Therefore, a detailed understanding of the functional connectivity between these nuclei and different brain regions is essential to gain insight into brain function and cognitive processes.

The functional and structural connectivity of thalamic nuclei has been investigated in numerous functional magnetic resonance imaging (fMRI) studies (Kumar et al., 2017, Mastropasqua et al., 2015, Kark et al., 2021) and DTI (Behrens et al., 2003, Jang et al., 2014, Lambert et al., 2017, Grodd et al., 2020). For example, resting-state fMRI (rs-fMRI) studies have revealed the existence of large-scale thalamocortical networks spanning many brain regions that involve multiple thalamic nuclei and their connections to different cortical regions (Kumar et al., 2017, Kumar et al., 2022, Zhang et al., 2008, Beckmann et al., 2005, Zhang et al., 2010). In addition to rs-fMRI studies, task-based fMRI (tb-fMRI) studies (Zhang et al., 2013, Jarrett et al., 2024, Wang, 2024, Rodriguez-Sabate et al., 2015) have played a crucial role in uncovering the functional interactions between thalamic nuclei, cortical, subcortical, and cerebellar regions during various cognitive and motor tasks. For example, Spets et al. (Spets and Slotnick, 2020) investigated the functional connectivity between anterior and mediodorsal thalamic nuclei with memory-related regions such as the prefrontal cortex, parietal cortex, visual processing regions, hippocampus, and parahippocampal cortex. Tb-fMRI studies have also elucidated the thalamocortical-cerebellar circuitry involved in motor control and coordination (Stoodley et al., 2012, Dacre et al., 2021).

Additionally, alterations in the functional connectivity of the thalamic nuclei have been observed in several neurological and psychiatric disorders, yet the causal relationship between these findings and the overarching disorders are not well understood. For example, disruptions in thalamocortical connectivity have been found to be implicated in progressive movement disorders, particularly Parkinson’s disease (Hao et al., 2020, Owens-Walton et al., 2019, Wang et al., 2021), emphasizing the importance of thalamocortical interactions in motor control. Altered thalamocortical connectivity has also been observed in cases of schizophrenia (Woodward et al., 2012, Wagner et al., 2015), depression (Sun et al., 2023, Brown et al., 2017), and attention-deficit/hyperactivity disorder (Hong, 2023, Kowalczyk et al., 2022), reflecting the impact on cognitive dysfunction.

Given the functional importance of the thalamic connectivity with different brain areas, the present study aims to investigate the functional connectivity of the thalamic nuclei during active motor (finger-tapping) and passive (tactile-finger) sensory tasks using high-resolution fMRI datasets acquired in our previous study (Charyasz et al., 2023). In this earlier study, we characterized functional localization and incidence of activation in thalamic nuclei involved in sensorimotor processes and assessed intersubject variability and reproducibility using tb-fMRI at 9.4T. Using GLM analysis, we identified distinct activation patterns within the group of lateral nuclei (VPL, VA, VLa, and VLp) and the group of pulvinar nuclei (PuA, PuM, and PuL) during both tasks. We also observed functional activation in the intralaminar nucleus group (CM and Pf) during the finger-tapping task. Starting from these findings, we now explore the functional connectivity between the identified thalamic nuclei and various cortical and subcortical areas as well as the cerebellum in this work. This may significantly enhance understanding of the complex functional interactions of thalamic nuclei involved in sensorimotor processing.

## 2. MATERIALS AND METHODS

### Participants

Our participant pool included eight healthy right-handed adults, with normal or corrected-to-normal vision, and an average age of 27 years (ranging from 21 to 34 years; five of whom were female). Ethical approval for this study was obtained from the local research ethics committee, and all participants provided written informed consent prior to participation.

### Experimental procedure

Each single session involved each participant to actively complete two fMRI block design tasks using their right hand: the tactile-finger task and the finger-tapping task. Please refer to our previous work (Charyasz et al., 2023) for a detailed description of the experimental setup and pre-processing pipeline used in this study.

### Tactile-finger task

The tactile-finger stimulation paradigm consisted of 12 cycles, with each cycle divided into alternating blocks of tactile stimulation (ON phase) and rest (OFF phase). Each block lasted a total of 40 seconds, equally divided into 20-second intervals for the ON and OFF phases, respectively. The tactile stimulation was delivered by air pulses through an inflatable finger clip, simultaneously applied to the thumb, index finger, middle finger, and ring finger. These air pulses, generated at a pressure of 2.5 bar, induced the displacement of the pneumatic membrane towards the skin surface for a duration of 250 milliseconds. To ensure a consistent and rhythmic pattern of stimulation across all fingers, the frequency of pulse delivery was set at 1 Hz. To prevent habituation and maintain participants’ attention, a random number of pulses (ranging from zero to four) was intentionally (deliberately) skipped within each stimulation block. This resulted in an average of 210-240 air pulses being delivered to each fingertip during each run. Participants were instructed to report the total number of blocks in which pulses were missing during the breaks between stimulation runs. Throughout the entire experiment, participants were lying still and keeping their eyes fixed on a black fixation cross on the screen.

### Finger-tapping task

The experimental design involved a visually-guided finger-tapping paradigm consisting of 12 visually cued cycles, each with a duration of 41 seconds. These cycles were divided into alternating blocks of finger tapping (ON phase) and rest (OFF phase) with each phase lasting 20 seconds. Each block of movement was preceded by a preparatory interval of 1 second, which was provided to ensure readiness. Participants were instructed to tap their right fingers sequentially, starting with the index finger, followed by the middle finger, ring finger, and little finger, against the thumb. The tapping rate was controlled by a visual cue in the form of a blinking arrow, with an approximate frequency of 2.5 Hz. During the designated rest blocks, participants were explicitly instructed to keep their eyes fixed on the black fixation cross, refraining from any voluntary movements.

### MR data acquisition

In a single session, both anatomical and functional images were acquired using a 9.4T whole-body MRI scanner (Siemens Healthineers, Erlangen, Germany) with an in-house-built head-coil equipped with 16 transmit and 31 receive channels (Shajan et al., 2014). In a separate session utilizing a Siemens Healthineers Prisma Fit 3T whole-body MRI scanner with a 64-channel head coil, high-resolution whole-brain anatomical images were acquired to facilitate the precise segmentation of the cortical and thalamic regions.

#### 9.4T imaging

High-resolution T1-weighted images were acquired using a magnetization-prepared rapid acquisition gradient echo (MPRAGE) sequence with the following parameters: inversion repetition time (TR) = 3.8 s, echo time (TE) = 2.50 ms, flip angle (FA) = 6°, field of view (FOV) = 192 mm, 288 sagittal slices covering the entire brain, voxel size = 0.6 x 0.6 x 0.6 mm3, GRAPPA acceleration factor (R) = 2 x 2, and partial Fourier in slab duration = 6/8. Task-based fMRI scans were collected using a 2D gradient-echo multi-band (MB) echo-planar imaging (GE-EPI) sequence with the following parameters: TR = 2 s; TE = 22 ms; FA = 50°; FOV = 198 mm; 86 interleaved slices per volume; voxel size = 1.25 x 1.25 x 1.25 mm^3^, *R* = 4, MB factor = 2, bandwidth = 1666 Hz/Px, and anterior-posterior phase encoding. For precise distortion correction, a set of ten volumes with reversed phase encoding (posterior-anterior) MB-GE-EPI scans was acquired, employing the exact same parameters as during the functional scans. Each subject underwent seven runs of the tactile-finger task (255 volumes, ~8.5 minutes per run), and a single run of the finger-tapping task (265 volumes, ~9 minutes).

#### 3T imaging

A comprehensive set of high-resolution T1-weighted and T2-weighted images were collected for each participant. The T1-weighted images were acquired using a magnetization-prepared rapid acquisition gradient echo (MPRAGE) sequence with TR = 2.4 s, TE = 2.22 ms, FA = 8°, FOV = 256 mm, and voxel size of 0.8 × 0.8 × 0.8 mm³. This acquisition yielded 208 sagittal slices. Additionally, the T2-weighted images were obtained using a 3D fast spin echo sequence with TR = 3.2 s, TE = 563 ms, FOV = 256 mm, and the same voxel size and slice number as the T1-weighted images.

### MRI data analysis

#### Data pre-processing

Both the task-based functional and structural images were preprocessed following the previously described methods (Charyasz et al., 2023). Briefly, the initial five volumes of the functional images were excluded, followed by correction for slice-time and head motion using the SPM12 software (R7771, http://www.fil.ion.ucl.ac.uk/spm). Furthermore, image distortions were corrected using the TopUp (Andersson et al., 2003) tool from the FSL package (Smith et al., 2004), while NORDIC (Moeller et al., 2021) denoising was employed to correct for thermal noise fluctuations. The resulting distortion-corrected datasets were subsequently co-registered with the anatomical data and spatially smoothed with a 2.5 mm full-width-at-half-maximum Gaussian kernel.

The 3T anatomical images, comprising T1-weighted and T2-weighted scans, were preprocessed with the FreeSurfer (http://surfer.nmr.mgh.harvard.edu, version 6.1) (Fischl et al., 1999) software package (version 6.1). The recon-all function within FreeSurfer was utilized to perform a thorough whole-brain segmentation. To accurately identify thalamic nuclei, a probabilistic thalamic segmentation algorithm (Iglesias et al., 2018), integrated within FreeSurfer, was employed. Additionally, the ACAPULCO (Kerestes et al., 2022) processing pipeline was applied specifically to the T1-weighted images for cerebellar segmentation. This involved utilizing the ACAPULCO tool to segment the cerebellum into lobules, enabling a detailed sub-segmentation of the cerebellar structure. The resulting segmentation outcomes served as regions of interest (ROIs) for further analysis.

#### Regions-of-interest (ROIs) selection

In accordance with the experimental design, which included tactile and motor tasks visually cued with attentional engagement, 66 anatomical ROIs (Table 1) were selected to investigate cortical-thalamic-cerebellar functional connectivity. Among these ROIs, 17 bilateral thalamic nuclei were chosen based on their prior identification (Charyasz et al., 2023) and their relevance to the research context. In addition, a set of 32 anatomical ROIs (16 in each hemisphere) comprising brain regions involved in sensorimotor, visual, and attentional signal processing were included. More specifically, 24 cortical ROIs were obtained from the Desikan-Killiany-Tourville (DKT) atlas, while the remaining 8 cerebellar ROIs were derived from the ACAPULCO segmentation.

**Table 1.** The complete list of the 33 bilateral ROIs and their abbreviations.

|  |  |
| --- | --- |
| ***Thalamic nuclei*** | anteroventral (AV), lateral dorsal (LD), lateral posterior (LP), ventral anterior (VA), ventral lateral anterior (VL<sub>a</sub>), ventral lateral posterior (VL<sub>p</sub>), ventral posterior lateral (VPL), centromedian (CM), parafascicular (Pf), lateral subdivision of mediodorsal thalamus (MD<sub>l</sub>), medial subdivision of mediodorsal thalamus (MD<sub>m</sub>), lateral geniculate (LGN), medial geniculate (MGN), anterior pulvinar (Pu<sub>A</sub>), inferior pulvinar (Pu<sub>I</sub>), later pulvinar (Pu<sub>L</sub>), medial pulvinar (Pu<sub>M</sub>) |
| ***Cortical regions*** | Paracentral gyrus (PCG), postcentral gyrus (PoCG), precentral gyrus (PrCG), superior parietal lobule (SPL), inferior parietal lobule (IPL), cuneus (Cu), pericalcarine (PCAL), lingual gyrus (LING), insula (INS) |
| ***Subcortical regions*** | Caudate (Ca) , Putamen (Pu) , Pallidum (Pd) |
| ***Cerebellum*** | Lobule I-II (CerLobI-III), Lobule IV (CerLobIV), Lobule V (CerLobV), Lobule VI (CerLobVI) |

#### Functional connectivity analysis

Task-based functional connectivity analyses were carried out using the CONN (Whitfield-Gabrieli and Nieto-Castanon, 2012) release 22a (Nieto-Castanon and Whitfield-Gabrieli, 2022) within MATLAB 2017b (The MathWorks, Inc., Natick, MA, USA). The preprocessed fMRI data were used as input to the CONN toolbox to undergo denoising and subsequent functional connectivity analysis. The functional data were denoised using a standard denoising pipeline (Nieto-Castanon, 2020). This denoising process included physiological noise correction achieved by regressing out 5 CompCor noise components from the white matter time series and 5 CompCor noise components from the CSF time series. In addition, motion regressors and their first-order derivatives (12 components) were used to account for any motion-related confounds. To disentangle task-related coactivations from genuine functional connectivity patterns, session and task effects along with their first-order derivatives (a total of 6 factors) were considered. Subsequently, linear detrending and band-pass filtering (0.008 Hz and 0.09 Hz) were applied on the BOLD timeseries (Hallquist et al., 2013) to reduce low- and high-frequency noise. CompCor (Behzadi et al., 2007, Chai et al., 2012) noise components within white matter and CSF were estimated by computing the average BOLD signal as well as the largest principal components orthogonal to the BOLD average within each subject’s eroded segmentation masks. From the number of noise terms included in this denoising strategy, the effective degrees of freedom of the BOLD signal after denoising were estimated to range from 652.7 to 667.5 (average 665.6) across all subjects (Nieto-Castanon, 2020).

Weighted ROI-to-ROI connectivity analysis (wRRC) was conducted to estimate functional connectivity matrices characterizing the functional interaction between each pair of 66 predefined ROIs. The strength of functional connectivity was represented by Fisher-transformed correlation coefficients from a weighted general linear model (GLM) (Nieto-Castanon, 2020b). For each pair of ROIs, coefficients were calculated independently to quantify the relationship between their respective BOLD signal time series. ROI time series were obtained for each participant by averaging the signal across all voxels within each ROI from pre-processed, unsmoothed fMRI data. Individual scans were weighted by a boxcar function representing the conditions of each task (tactile and motor), convolved, and rectified with a canonical SPM hemodynamic response function. Second-level analyses were conducted using a GLM approach (Nieto-Castanon, 2020c). First-level connectivity measures served as dependent variables, with one independent sample per participant and one measure per task to estimate a separate GLM for each connection. Connection-level hypotheses were tested using multivariate parametric statistics with random effects modeling across participants and covariance estimation across measures. Statistical inference was performed at the level of individual functional connections, and results were thresholded using a family-corrected p-FDR < 0.05 connection-level threshold (Benjamini et al., 2001).

## 3. RESULTS

The results of our ROI-based functional connectivity analysis clearly showed task-dependent changes in inter-regional functional connectivity between the selected ROIs. These changes were dependent on whether participants performed a finger-tapping task or a tactile task. To capture the overall connectivity patterns, the connectivity matrices showing the task-modulated interactions between all 66 ROIs across all subjects are presented in Figure 1, providing a comprehensive representation of the connections. In general, robust connections were observed between and among thalamic, cortical, subcortical and cerebellar regions in both hemispheres. We also divided the data into smaller groups for cross-comparison intended to improve clarity and organization while isolating task-specific changes in connectivity within the examined brain regions. It is important to note that the results of task-dependent functional connectivity between cortical, subcortical, and cerebellar areas are not included in this section. These results are presented for future reference in the Supplementary section (Figure S1 to Figure S4).

**Figure 1.**
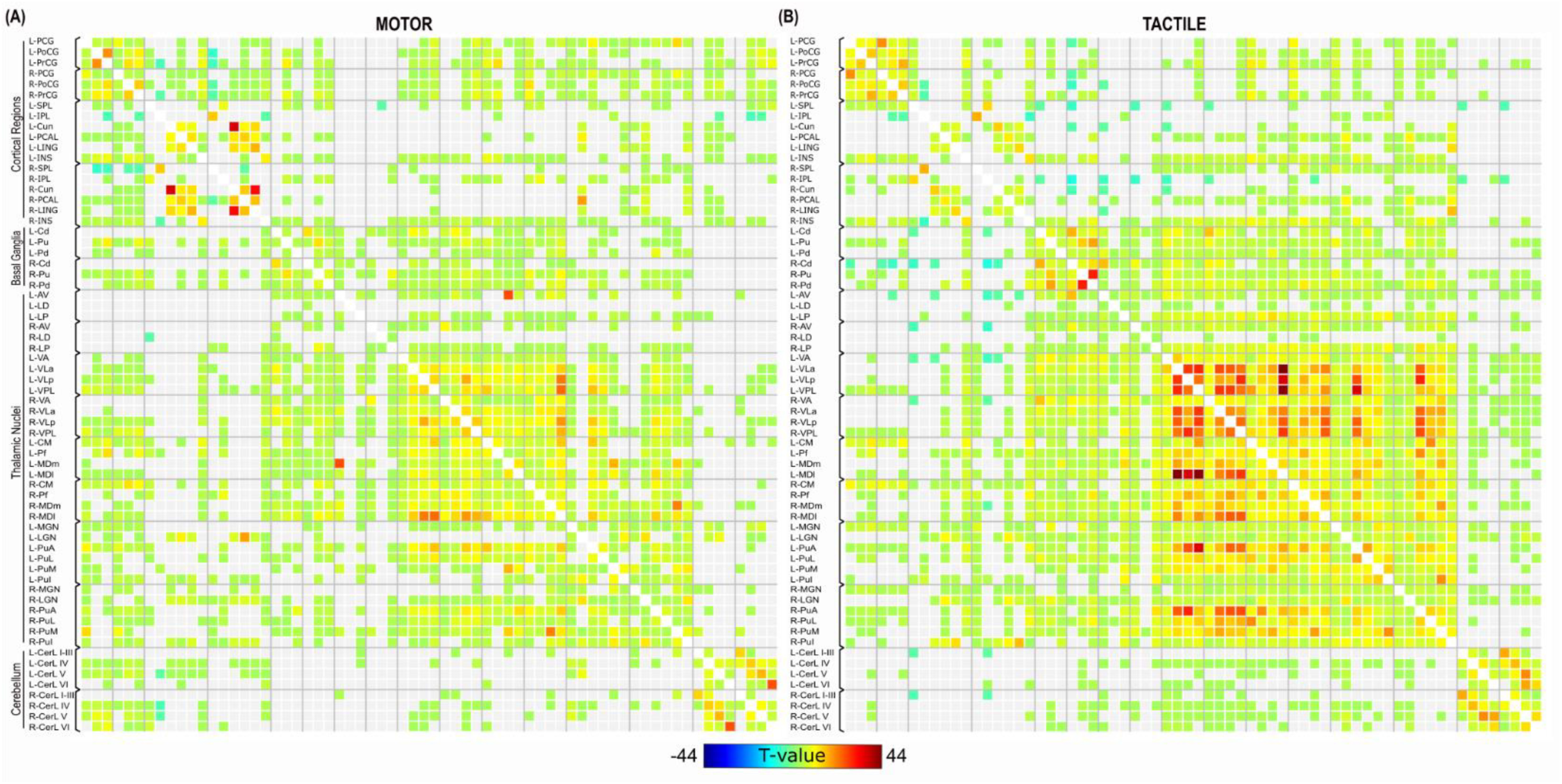
Average ROI-to-ROI functional connectivity matrices for the motor (A) and tactile (B) tasks across 66 ROIs across subjects, thresholded at corrected p-FDR < 0.05 connection level threshold.

### 3.1 Task-dependent functional connectivity of the anterior thalamic nuclei

In both motor and tactile tasks, the connectivity analysis revealed increased functional connectivity between the anterior thalamic nuclei (AV, LD and LP) and various brain regions, including cerebellar lobules and basal ganglia structures (caudate, putamen and pallidum). In the motor task, increased connectivity was observed between the right LP nucleus and the cortical regions of the right insula, right SPL, and right IPL. Decreased connectivity was found between the left SPL and the right LD nucleus. In the tactile task, increased connectivity was observed between bilateral AV and LP nuclei and the bilateral insula, PCAL and IPL. In addition, decreased connectivity was found between AV nuclei and bilateral cuneus, right lingual gyrus, bilateral SPL and right PoCG. In general, the tactile task exhibited a higher number of significant connections compared to the motor task. The comprehensive details of these observed connections, including the specific brain regions involved and the corresponding t-values, are summarized in Supplementary Table S1. In addition, the graphical representation of these connectivity patterns is shown in Figure 2.

**Figure 2.**
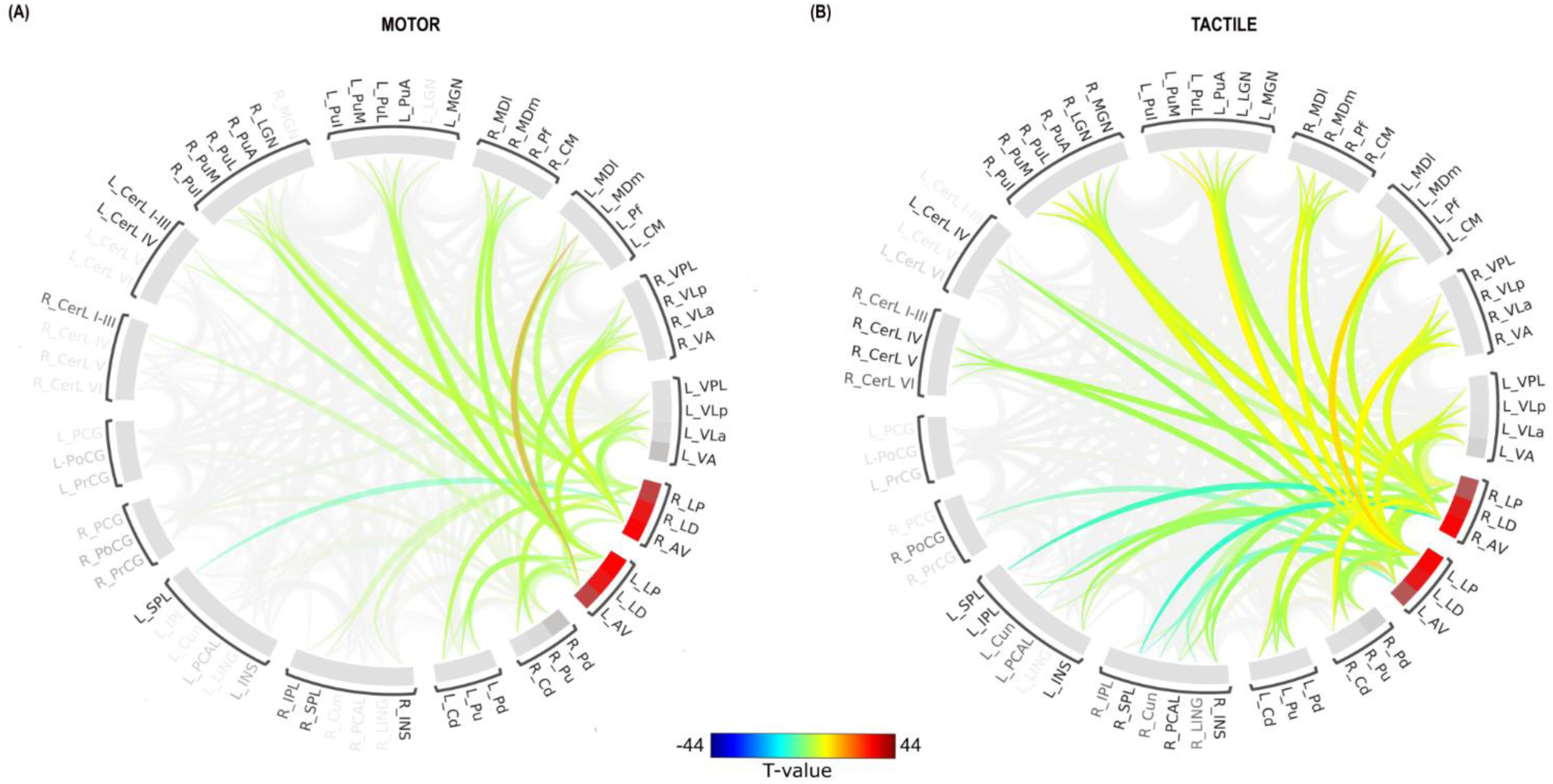
ROI-to-ROI connectome ring of functional connectivity between anterior thalamic nuclei and other brain regions for the motor (A) and tactile (B) tasks. The color links represent ROI-to-ROI connections with a false discovery rate (FDR) of p < 0.05. Warm colors (yellow and red) indicate increased connectivity, while cool colors (dark and light blue) signify decreased connectivity.

### 3.2 Task-dependent functional connectivity of the medial thalamic nuclei

Analysis of the connectivity between the medial thalamic nuclei (CM, Pf, MDm and MDl) and other brain regions revealed distinct variations in the connectivity patterns between the motor and tactile tasks are shown in Figure 3. The detailed Supplementary Table S2 provides a comprehensive overview of all significant connections and their corresponding statistical values.

**Figure 3.**
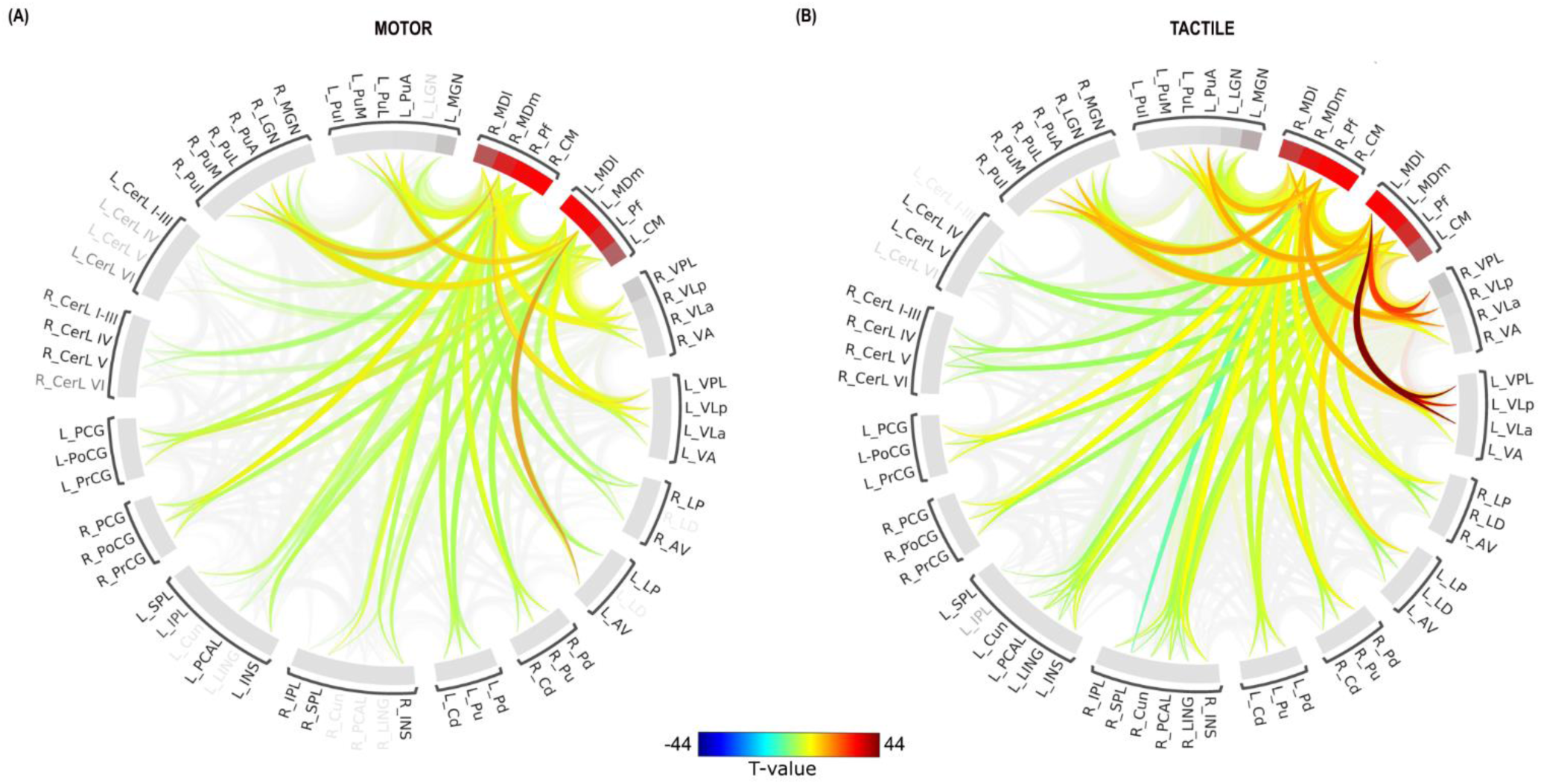
ROI-to-ROI connectome ring of functional connectivity between medial thalamic nuclei and other brain regions for the motor (A) and tactile (B) tasks. The color links represent ROI-to-ROI connections with a false discovery rate (FDR) of p < 0.05. Warm colors (yellow and red) indicate increased connectivity, while cool colors (dark and light blue) signify decreased connectivity.

Increased connectivity between cerebellar lobules and thalamic nuclei was observed in both motor and tactile tasks, with notable differences in the specific nuclei involved. In the motor task, connectivity was predominantly observed with the CM and MDm nuclei, whereas in the tactile task all four nuclei of the medial thalamic group showed increased connectivity. The number of significant connections was notably lower in the motor task than in the tactile task. In both tasks, there was a significant increase in functional connectivity between all four nuclei of the medial group and bilateral basal ganglia structures, including the putamen, pallidum and caudate, with a similar number of connections.

The medial group nuclei connectivity patterns differed between the two tasks. Specifically, connectivity between two nuclei (CM and PF) and both bilateral cunei and bilateral lingual areas was observed in the tactile task, whereas no such connectivity was observed in the motor task. There were also significant differences in connectivity to the PCAL region. In the tactile task, increased connectivity was observed between all four nuclei and bilateral PCAL. In contrast, only two connections were found in the motor task, specifically between the bilateral CM nuclei and the left PCAL.

Interestingly, the most pronounced differences in connectivity patterns were found in the sensorimotor cortical regions, including the PCG, PoCG and PrCG. Both tasks resulted in connections between bilateral PCG and bilateral CM, albeit a greater number of connections were observed in the motor task. The bilateral MDL and MDm, however, only resulted in statistically significant connections during the motor task (not the tactile task). In the motor task, the bilateral PoCG showed significant connections with all four medial thalamic nuclei, whereas the bilateral PrCG showed significant connections with three thalamic nuclei (CM, Pf and MDl). Similarly, during the tactile task, the bilateral PoCG and PrCG exhibited significant connections with all four nuclei.

### 3.3 Task-dependent functional connectivity of the lateral thalamic nuclei

Examining the connectivity between the lateral thalamic nuclei (VA, VLa, VLp and VPL) and other brain regions revealed interesting variations in the connectivity patterns across the motor and tactile tasks, as shown in Figure 4. In this section, we provide a comprehensive overview of the observed connectivity patterns in each task, together with the corresponding statistical values, as summarized in Supplementary Table S3.

**Figure 4.**
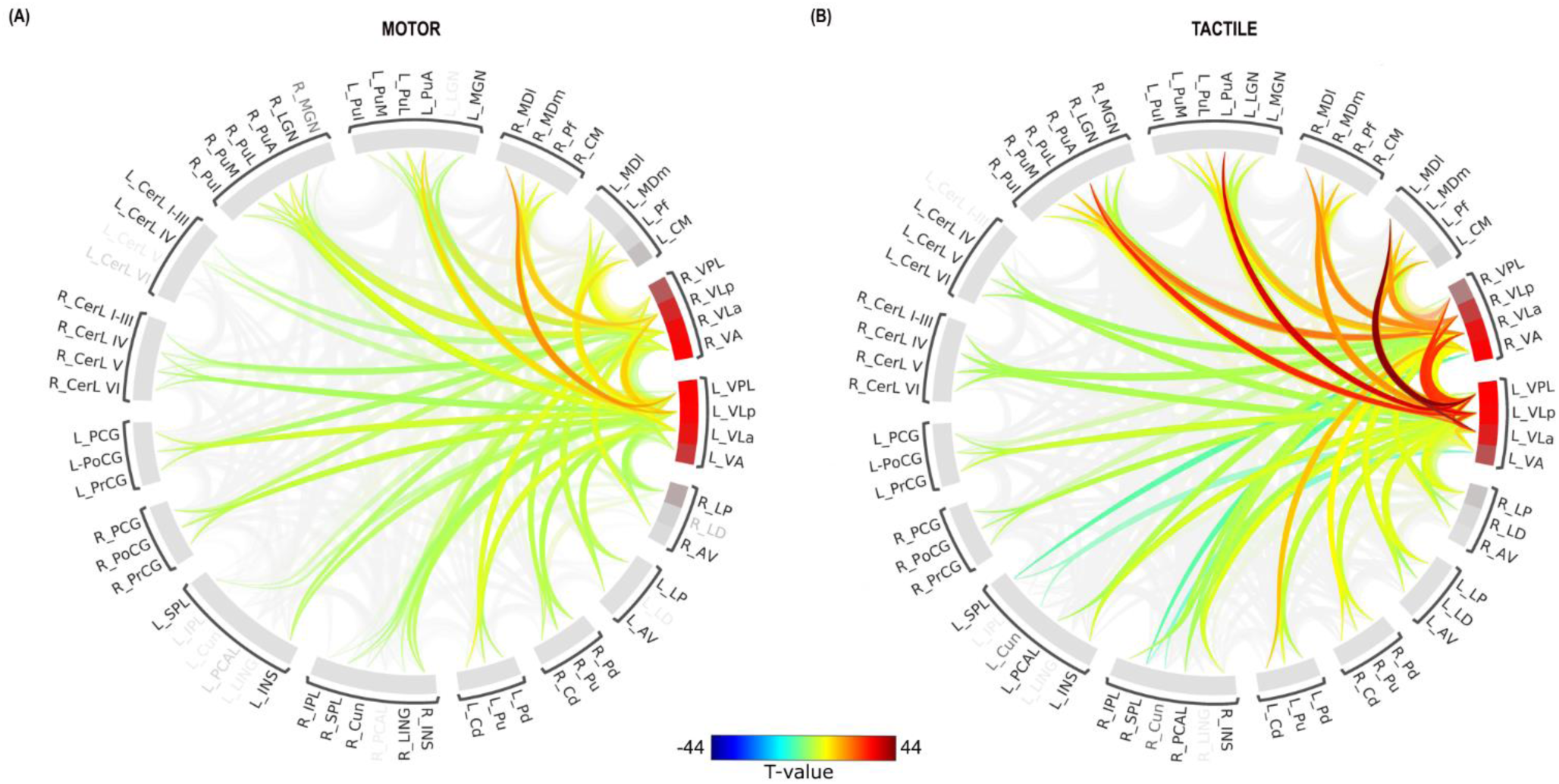
ROI-to-ROI connectome ring of functional connectivity between lateral thalamic nuclei and other brain regions for the motor (A) and tactile (B) tasks. The color links represent ROI-to-ROI connections with a false discovery rate (FDR) of p < 0.05. Warm colors (yellow and red) indicate increased connectivity, while cool colors (dark and light blue) signify decreased connectivity.

Consistent with the groups of anterior and medial thalamic nuclei, the motor and sensory tasks showed similar patterns of increased connectivity between the cerebellar lobes and the lateral thalamic nuclei. Notably, with fewer significant connections in the motor task, the strength of the connections was similar in both tasks. Both motor and tactile tasks showed increased connectivity between subcortical regions (bilateral putamen, caudate and pallidum) and all four bilateral nuclei (VA, VLa, VLp and VPL). However, compared to the motor task, the tactile task exhibited significantly higher connectivity strength. The bilateral insula showed increased connectivity with the bilateral nuclei of the lateral nuclear group in both motor and tactile tasks. The strength of these connections, however, was comparable between the tasks.

Increased connectivity between the bilateral PCAL and the bilateral VLa, VLP and VPL nuclei was only found during the tactile task, whereas the increased connectivity between the left VPL nucleus and the right lingual gyrus was only observed during the motor task. In the tactile task, the right IPL showed increased connectivity with all four nuclei bilaterally, whereas in the motor task connectivity was limited to a single connection with the left VLa nucleus. On the other hand, connectivity between the left VPL nucleus and the right cuneus increased during the motor task. In addition, several negative connections appeared only during the tactile task. These included a reduction in connectivity between the left and right SPL and the bilateral VA nucleus, and between the bilateral cuneus and the left VA nucleus.

Similar to the medial group of thalamic nuclei, the most significant differences in connectivity patterns between tasks were found between the lateral group of thalamic nuclei and sensorimotor cortical regions (PCG, PoCG and PrCG). Connectivity analysis revealed differences between the motor and tactile tasks in the number of pairwise connections. A total of 33 and 14 pairwise connections were observed in the motor task and the tactile task, respectively. In the motor task, increased connectivity was observed between the bilateral PCG and the VLa, VLp and VPL nuclei, with the strongest connections observed with the bilateral VPL. The PoCG reflected increased connectivity between the bilateral and all four nuclei. PrCG reflected enhanced connectivity bilaterally with the VLa, VLp, and VPL nuclei. Alternatively, the tactile task had different connectivity patterns. The tactile task reflected increased connectivity between the PCG, PrCG, and PoCG bilaterally with the VPL nucleus. The left PrCG also showed increased connectivity with the left VA and left VLp nuclei in the tactile task. Overall, the motor task showed a higher number and strength of connections with thalamic nuclei, particularly involving Vla, VLp, and VPL, compared to the tactile task, which involved mainly the VPL nucleus with fewer connections.

### 3.4 Task-dependent functional connectivity of the posterior thalamic nuclei

Consistent with findings in other thalamic nuclei groups, analysis of the posterior group (MGN, LGN, PuA, PUL, PuM, and PuI) revealed a significantly lower number of pairwise connections with subcortical and cerebellar regions in the motor task compared to the tactile task. Increased connectivity between the cerebellar lobules and the thalamic nuclei was observed in both the motor task and the tactile task, with notable differences in the specific nuclei involved. Enhancement of connectivity between the posterior group of nuclei (left LGN, bilateral MGN, bilateral PuA and PuM) and bilateral cerebellar lobules (I-III, IV and V) was observed in the motor task, whereas connectivity between bilateral nuclei (LGN, MGN, PuA, PuL and PuM) and bilateral lobules IV-VI was found in the tactile task. In both tasks, a significant increase in functional connectivity was observed between all nuclei of the posterior group and the bilateral nuclei of the basal ganglia, including the putamen, pallidum and caudate, with similar strength of connections.

Increased connectivity of thalamic nuclei with specific cortical regions was found in both tasks. Connectivity analysis of the tactile task revealed a greater strength and number of connections in the bilateral insula, whereas the bilateral SPL showed a greater number of connections in the motor task. The bilateral PCAL showed a greater strength of connectivity in the motor task but, interestingly, with a greater number of connections in the tactile task. The cuneus and the lingual gyrus showed consistent connectivity patterns bilaterally in both tasks. Notably, the bilateral IPL only showed increased connectivity in the tactile task.

Similar to the previous thalamic nuclei groups, differences between tasks in the pattern of connectivity with sensorimotor cortical regions were revealed. An increase in connectivity between the distinct nuclei of the lateral group and the bilateral PCG, PoCG and PrCG was observed in both tasks. However, the number of pairwise connections was higher in the motor task (46 connections) than in the tactile task (37 connections). In the motor task, the bilateral PCG showed stronger connections with all nuclei belonging to the lateral group. However, in the tactile task, no connectivity was observed between the PCG and LGN and PuM nuclei. Both tasks showed comparable connectivity strength between the PoCG and other nuclei, with no connections to LGN nuclei in either task. Similarly, the bilateral PrCG showed no connectivity with PuM nuclei in either task, and with no PuI in the tactile task.

Supplementary Table S4, which provides detailed information on all detected pairwise connections, shows these task-dependent differences in connectivity. Furthermore, Figure 5 shows them visually, where panel A shows the motor task and panel B shows the tactile task.

**Figure 5.**
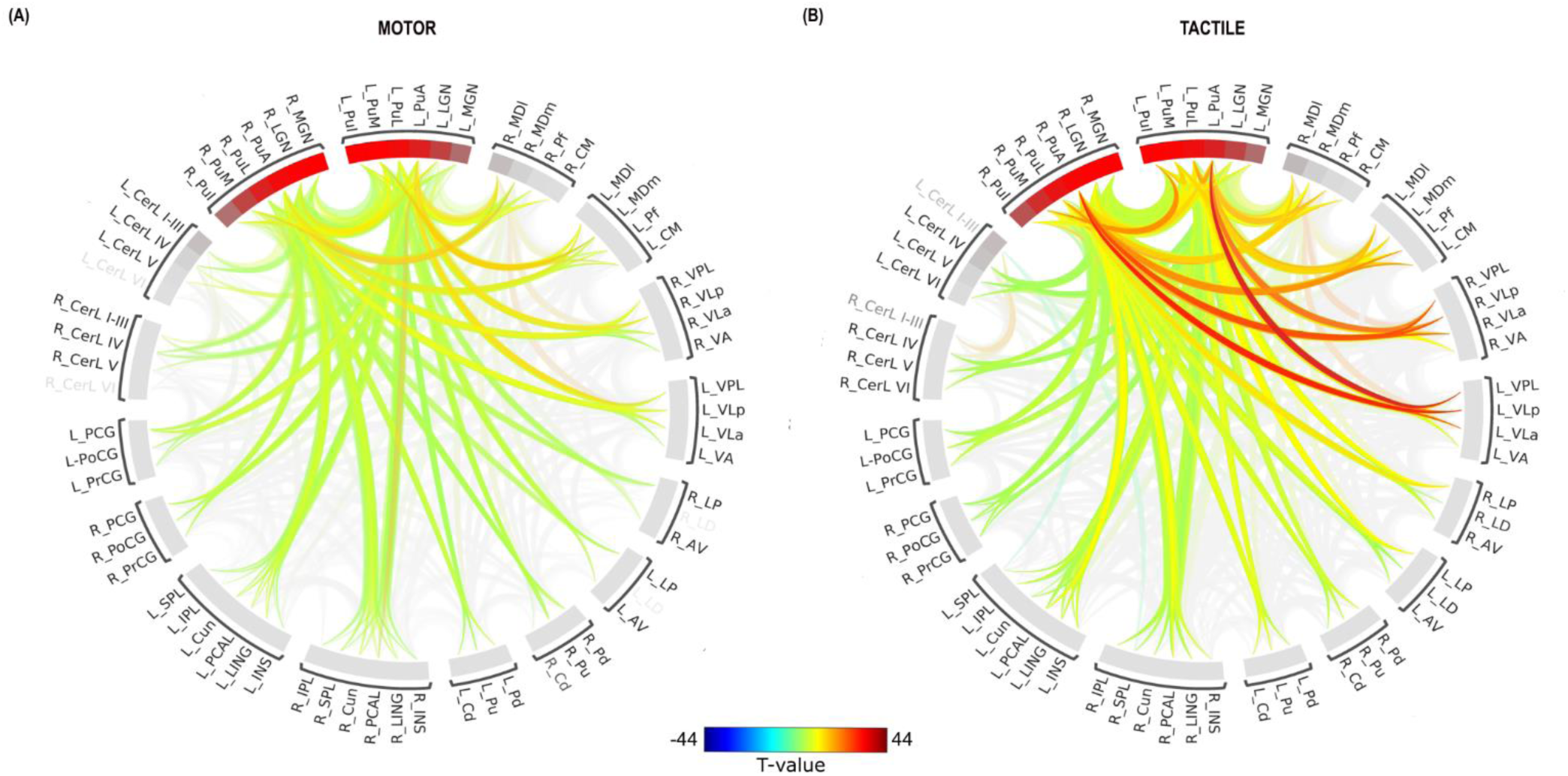
ROI-to-ROI connectome ring of functional connectivity between posterior thalamic nuclei and other brain regions for the motor (A) and tactile (B) tasks. The color links represent ROI-to-ROI connections with a false discovery rate (FDR) of p < 0.05. Warm colors (yellow and red) indicate increased connectivity, while cool colors (dark and light blue) signify decreased connectivity.

## 4. DISCUSSION

The aim of this study was to extend our previous work (Charyasz et al., 2023) by conducting a detailed examination of task-dependent functional connectivity between thalamic nuclei and various cortical, subcortical, and cerebellar regions. While the results of the voxel-based analysis in our previous study identified the specific thalamic nuclei activated during both tasks, they did not provide any insight into the connections between these nuclei and different brain regions. To address this limitation, we analyzed our previously published dataset of task-based fMRI measurements at 9.4T with a focus on the assessment of connectivity patterns during both an active motor (finger-tapping) task and a passive (tactile-finger) sensory task.

### Task-Dependent and Region-Specific Functional Connectivity

The identification of both task-dependent as well as region-specific variability in the functional connectivity was an important part of our study. Specifically, the tactile task exhibited a higher number of significant connections compared to the motor task. The stronger connectivity of the thalamic nuclei with regions such as insula, IPL and PCAL suggests that sensory integration and attentional processing are enhanced during passive stimulation. These results are coherent with previous work suggesting that the thalamus does not simply act as a passive relay station but plays an active role in adapting sensory processing (Sherman, 2016, Hwang et al., 2017). Furthermore, the involvement of visual regions, including the cuneus and lingual gyrus, in the tactile task highlights the multisensory nature of thalamic processing (Barczak et al., 2023, Driver and Noesselt, 2008). In contrast, consistent with the role of the thalamus in motor coordination and proprioception (Bosch-Bouju et al., 2013, Semrau et al., 2015), the motor task showed more focused connectivity with cerebellar and cortical motor areas. The observed differences in strength and connectivity patterns between the two tasks show the ability of the thalamus to dynamically respond to the specific functional demands of different sensorimotor tasks. A more detailed discussion of how the nature of the task and inclusion of thalamic groups contribute to these connectivity patterns is provided below.

### Anterior thalamic nuclei (ATN)

The anterior thalamic nuclei - AV, LD and LP - are known for their role in memory, learning and attention. These nuclei are essential components of the Papez circuit (Papez, 1995), which connects the thalamus to the hippocampus and cingulate cortex to support cognitive and emotional functions (Jankowski et al., 2013, Grodd et al., 2020, Aggleton et al., 2010, Nelson, 2021). In addition to these functions, the ATN has also been implicated in integration, particularly in tasks requiring focused attention and sensory discrimination (Wright et al., 2015, Sweeney-Reed et al., 2017). Enhanced connectivity between the ATN and subcortical regions, including the cerebellar lobules and the basal ganglia (caudate, putamen, pallidum), was observed in both motor and tactile tasks, however with a higher number of pairwise connections in the tactile task. The direct involvement of the ATN in motor processes is not well-studied, but the connectivity observed here suggests a possible modulatory role in thalamo-striatal and thalamo-cerebellar pathways by supporting attention-driven sensorimotor integration.

The connectivity patterns specific to the two tasks show the versatility of the ATN. The motor task elicited increased connectivity of the ATN with the insula and IPL, which are associated with sensorimotor integration and movement planning (Chang et al., 2013, Mehler and Reschechtko, 2018). In contrast, the tactile task engaged a broader network, including the insula, the IPL, and the PCAL. This pattern of connectivity indicates a role for the ATN in multisensory integration and spatial representation (Jankowski et al., 2013). Furthermore, the observed reduction in connectivity with visual regions, such as the cuneus, can be interpreted as a process of neural reorganization that prioritizes tactile processing over visual input. This is consistent with the concept of flexible modulation within thalamocortical networks, which allows for task-specific reorganization depending on sensory demands (Sasaki et al., 2022).

### Medial thalamic nuclei

The medial thalamic nuclei (CM, Pf, MDm and MDl) are thought to act as important hubs in the thalamo-cortical and thalamo-striatal networks, which are involved in sensorimotor coordination, attention and higher-order cognitive functions (Jones, 2007, Kumar et al., 2023, Van der Werf et al., 2002). The tactile task demonstrates connectivity among all four medial thalamic nuclei, which is consistent with another study reporting that tactile processing involves a broader network of thalamic structures, demonstrating the ability of the thalamus to integrate multimodal sensory inputs (Habig et al., 2023, Wahlbom et al., 2021). The extensive involvement of thalamic nuclei in tactile tasks may enhance the complex sensory discrimination necessary for tactile signal processing exploration. The observed increase in connectivity between the medial thalamic nuclei and the basal ganglia (putamen, pallidum, and caudate) in both tasks supports previous findings that thalamo-striatal pathways play a role in motor and sensory signal processing (Smith et al., 2009). The consistent connectivity noted in both tasks shows that the thalamo-striatal network is not limited to specific tasks but may instead serve as an essential network for sensory integration.

Task-specific connectivity between medial thalamic nuclei and cortical regions, particularly the posterior cortical regions (cuneus, lingual gyrus, and PCAL), and the sensorimotor cortex (PCG, PoCG, and PrCG), also exhibit differences. While no connectivity was detected between the CM and Pf nuclei and the cuneus or lingual areas during the motor task, robust connectivity was evident during the tactile task. This finding is in line with those of previous studies showing that the lingual and the cuneus regions are involved in sensory processing and visual-spatial attention (Lee et al., 2020, Palejwala et al., 2021). An increased number of connections were found between sensorimotor cortex regions and the medial thalamic nuclei during the motor task, which aligns with the role of these nuclei in sensorimotor coordination, learning and decision making (Ilyas et al., 2019, Mitchell, 2015, Saalmann, 2014).

### Lateral thalamic nuclei

The lateral thalamic nuclei - VA, VL (VLa, VLp) and VPL - are referred to as the motor thalamic nuclei (Ilinsky et al., 2018) and play a crucial role in sensorimotor processing by acting as a hub for integrating and relaying sensory and motor information to various cortical and subcortical regions (Bosch-Bouju et al., 2013, Jones, 2007).

The enhanced connectivity between the lateral thalamic nuclei and cerebellar regions emphasizes the importance of the cerebello-thalamic pathway in motor coordination and sensory integration (Palesi et al., 2015, Pisano et al., 2021). In the motor task, increased connectivity was observed between the VLp, VPL, and cerebellar lobules (I-III, IV, V), aligning with the cerebellum’s role in delivering motor feedback and error correction during movement (Stoodley et al., 2012, Popa et al., 2016). Cerebellar outputs project to the VA and VL nuclei to augment motor execution, further supporting the role of the thalamus as a dynamic relay and control hub in motor signal processing (Gornati et al., 2018, Koster and Sherman, 2024). In the tactile task, increased connectivity was observed in the cerebellar lobules IV-VI with stronger involvement of the VL nuclei, indicating an involvement for these thalamic nuclei in the integration of tactile sensory input with motor adjustments.

Connectivity between the lateral thalamic nuclei and the basal ganglia (putamen, caudate, and pallidum) confirms their importance in human motor and sensory circuits. The VA, VL and VPL nuclei show increased connectivity with the putamen and pallidum in the motor task, suggesting their involvement in the cortico-basal ganglia-thalamo-cortical loop. Evidence from human studies corroborates this circuit, demonstrating that basal ganglia outputs modulate thalamic activity to ensure motor execution (Draganski et al., 2008, Mohagheghi Nejad et al., 2018, McFarland and Haber, 2002). Enhanced connectivity with subcortical regions in the tactile task indicates a greater demand for sensory integration and processing, consistent with the concept of task-specific recruitment of neural resources (Jang et al., 2017, Sasaki et al., 2022).

The lateral thalamic nuclei exhibit a higher number of significant connections with sensorimotor cortical regions during the motor task compared to the tactile task. This variability suggests that the thalamic nuclei may have distinct roles in modulating or integrating signals based on the complexity and demands of tasks (Wang, 2024, Halassa and Kastner, 2017). The PrCG consistently showed connection with the VA, VLa, VLp, and VPL nuclei across tasks, supporting the integration of motor-related signals with sensory feedback for movement planning and execution. This is in line with research that shows how the VL and VPL nuclei relay information from the cerebellum and basal ganglia to cortical motor regions, ensuring coordinated and precise movement (Middleton and Strick, 2000, Sommer, 2003). The PoCG exhibited consistent connection with the bilateral VPL nuclei in both tasks, emphasizing the VPL’s role in relaying tactile information. This finding aligns with research demonstrating that the VPL supports sensory integration by relaying inputs from ascending pathways to cortical regions (Studtmann et al., 2023). Decreased connectivity between the VA nucleus and bilateral cuneus, as well as the bilateral SPL may be provoked by the active suppression of task-irrelevant regions (Tomasi et al., 2014).

### Posterior thalamic nuclei

The posterior thalamic nuclei - LGN, MGN, PuA, and PuM – play a crucial role in sensory and cognitive processing by integrating and transmitting visual, auditory, and multisensory information, as well as supporting attention, sensory prioritization, and cognitive control (Saalmann et al., 2012, Saalmann and Kastner, 2011, Nagalski et al., 2016, Meng and Schneider, 2022). These nuclei show distinct patterns of connectivity with cerebellar, subcortical, and cortical regions, reflecting their adaptive roles in sensory and motor processing. In the motor task, increased connectivity was observed in the LGN, MGN, PuA, and PuM and cerebellar lobules I-III, IV, and V. The LGN’s involvement is consistent with its role in relaying visual information essential for visuomotor coordination (Casagrande et al., 2005, Lesica and Stanley, 2005). Similarly, the MGN’s connections might reflect the importance of auditory-motor integration for tasks requiring timing and rhythm (O’Connor et al., 1997). The pulvinar nuclei (PuA and PuM) exhibit functional connectivity with these cerebellar lobules, supporting visuomotor coordination and attentional control, aligning with their known role in modulating sensorimotor processing (Bridge et al., 2016, Froesel et al., 2021, Saalmann and Kastner, 2011). The tactile task elicits connectivity with cerebellar lobules IV-VI, with the LGN nuclei exhibiting stronger interactions, suggesting their involvement in tactile sensory processing and attentional modulation. Thalamic connectivity with basal ganglia regions, including the putamen, pallidum, and caudate, remains consistent across tasks. In both tasks, all posterior group nuclei exhibit strong connectivity with basal ganglia regions, suggesting their involvement in voluntary movements and visual signal processing (Wilke et al., 2018, Cortes et al., 2024, Shimono et al., 2012).

Thalamocortical connectivity patterns also differed between tasks. In the motor task, the LGN, MGN, and pulvinar nuclei show connections with sensorimotor regions, including the PrCG, PoCG, PCG and SPL. These connections highlight their role in motor execution, sensory feedback integration, and visuospatial processing, consistent with research on the role of the thalamus in coordinating sensorimotor pathways (Basile et al., 2024). In the tactile task, the pulvinar nuclei exhibit enhanced connectivity with the insula and IPL, which are regions associated with tactile information processing and attentional regulation (Barron et al., 2015, Homman-Ludiye and Bourne, 2019).

### Summary of findings

Our results reveal distinct task-specific functional connectivity patterns in the anterior, medial, lateral, and posterior thalamic nuclei. Instead of acting as static hubs with fixed roles in neural signal processing, these nuclei function as versatile hubs that dynamically adjust their functional connections in response to the demands of different tasks. During the motor task, thalamic networks were primarily involved in motor planning, execution, and proprioceptive feedback, whereas the tactile task elicited broader connectivity with regions associated with sensory integration, attentional control, and visual processing. This confirms and extends previous observations of the heterogeneity of thalamic nuclei in sensorimotor processing (Kumar et al., 2022, Zhou et al., 2016, Saalmann and Kastner, 2015, Acsady, 2023).

### Limitations and future directions

Although this study provides valuable insights, certain limitations must be acknowledged. The study relies on the relatively small sample size, the use of stimuli applied only to the right hand, and the challenges related to accurate segmentation of thalamic nuclei due to altered tissue contrast at 9.4T as compared to lower clinical field strengths. Nevertheless, the dataset provided sufficient spatial resolution and signal quality to investigate task-dependent activation and connectivity patterns and provide robust insights into the functional role of the thalamic nuclei. The understanding of thalamic connectivity could be further improved by addressing these limitations in future research. Increasing the sample size, applying stimulation to both hands and using improved segmentation techniques may provide a more comprehensive view of the thalamic function. An extended range of performed sensorimotor tasks would also help to confirm and extend these findings. Furthermore, investigating thalamic connectivity in clinical populations, such as Parkinson’s disease patients (Halliday, 2009, Wang et al., 2021), could provide important information on how connectivity alterations affect sensorimotor impairments.

To the best of our knowledge, this study is the first to show comprehensive results of task-dependent functional connectivity between thalamic nuclei and cortical, subcortical, and cerebellar regions using task-based fMRI at a field strength of 9.4T, which indicates the role of the thalamus as a flexible hub for sensorimotor integration. This study extends our current understanding of thalamic functional heterogeneity by demonstrating motor and tactile connectivity patterns. Mapping these dynamic connectivity patterns has important implications for future research on thalamic function in healthy and clinical populations.

## Abbreviations

AC: anterior commissure
AV: anteroventral
ATN: anterior thalamic nuclei
BOLD: blood-oxygen-level-dependent
CM: centromedian
Cun: cuneus
EPI: echo planar imaging
FA: flip angle
fMRI: functional magnetic resonance imaging
FOV: field of view
FWHM: full width at half maximum
GLM: general linear model
HRF: hemodynamic response function
INS: insula
IPL: inferior parietal lobule
LD: lateral dorsal
LGN: lateral geniculate
LING: lingual gyrus
LP: lateral posterior
MDl: lateral subdivision of mediodorsal thalamus
MDm: medial subdivision of mediodorsal thalamus
MGN: medial geniculate
MPRAGE: magnetization-prepared rapid acquisition gradient echo
PCG: paracentral gyrus
PCAL: pericalcarine
PC: posterior commissure
Pf: parafascicular
PoCG: postcentral gyrus
PuA: anterior pulvinar
PuI: inferior pulvinar
PuL: later pulvinar
PuM: medial pulvinar
PrCG: precentral gyrus
GRAPPA: acceleration factor
SPL: superior parietal lobule
TE: echo time
TR: repetition time
VA: ventral anterior
VL: ventral lateral
VLa: ventral lateral anterior
VLp: ventral lateral posterior
VP: ventral posterior
VPL: ventral posterior lateral

## Author contributions

EC: study design, stimulus programming, data acquisition, planning and performing the data analysis, writing–original draft, and preparing the figures. ME: advice for data evaluation and reviewing the manuscript. JB: advice for MR measurements and protocol optimization, reviewing and editing the manuscript. RH: assistance in data acquisition, reviewing and editing the manuscript. BB: reviewing and editing the manuscript. VK: reviewing and editing the manuscript. WG: supervision, reviewing and editing the manuscript. KS: resources, supervision, funding acquisition, reviewing and editing the manuscript. All authors contributed to the article and approved the submitted version.

## References

1. Acsady, L. 2023. 32Heterogeneity of Thalamic Input Landscapes. In: Bickford, M., Usrey, W. M. & Sherman, S. M. (eds.) The Cerebral Cortex and Thalamus. Oxford University Press.

2. Aggleton, J. P., O’mara, S. M., Vann, S. D., Wright, N. F., Tsanov, M. & Erichsen, J. T. 2010. Hippocampal-anterior thalamic pathways for memory: uncovering a network of direct and indirect actions. Eur J Neurosci, 31, 2292–307.

3. Andersson, J. L., Skare, S. & Ashburner, J. 2003. How to correct susceptibility distortions in spin-echo echo-planar images: application to diffusion tensor imaging. Neuroimage, 20, 870–88.

4. Azevedo, F. A., Carvalho, L. R., Grinberg, L. T., Farfel, J. M., Ferretti, R. E., Leite, R. E., Jacob Filho, W., Lent, R. & Herculano-Houzel, S. 2009. Equal numbers of neuronal and nonneuronal cells make the human brain an isometrically scaled-up primate brain. J Comp Neurol, 513, 532–41.

5. Barczak, A., O’connell, M. N. & Schroeder, C. E. 2023. 305Thalamic Contributions to Multisensory Convergence and Processing. In: King, A. J., Hirsch, J. A., Usrey, W. M. & Sherman, S. M. (eds.) The Cerebral Cortex and Thalamus. Oxford University Press.

6. Barron, D. S., Eickhoff, S. B., Clos, M. & Fox, P. T. 2015. Human pulvinar functional organization and connectivity. Hum Brain Mapp, 36, 2417–31.

7. Basile, G. A., Quartarone, A., Cerasa, A., Ielo, A., Bonanno, L., Bertino, S., Rizzo, G., Milardi, D., Anastasi, G. P., Saranathan, M. & Cacciola, A. 2024. Track-Weighted Dynamic Functional Connectivity Profiles and Topographic Organization of the Human Pulvinar. Hum Brain Mapp, 45, e70062.

8. Beckmann, C. F., Deluca, M., Devlin, J. T. & Smith, S. M. 2005. Investigations into resting-state connectivity using independent component analysis. Philos Trans R Soc Lond B Biol Sci, 360, 1001–13.

9. Behrens, T. E., Johansen-Berg, H., Woolrich, M. W., Smith, S. M., Wheeler-Kingshott, C. A., Boulby, P. A., Barker, G. J., Sillery, E. L., Sheehan, K., Ciccarelli, O., Thompson, A. J., Brady, J. M. & Matthews, P. M. 2003. Non-invasive mapping of connections between human thalamus and cortex using diffusion imaging. Nat Neurosci, 6, 750–7.

10. Behzadi, Y., Restom, K., Liau, J. & Liu, T. T. 2007. A component based noise correction method (CompCor) for BOLD and perfusion based fMRI. Neuroimage, 37, 90–101.

11. Benjamini, Y., Drai, D., Elmer, G., Kafkafi, N. & Golani, I. 2001. Controlling the false discovery rate in behavior genetics research. Behav Brain Res, 125, 279–84.

12. Bosch-Bouju, C., Hyland, B. I. & Parr-Brownlie, L. C. 2013. Motor thalamus integration of cortical, cerebellar and basal ganglia information: implications for normal and parkinsonian conditions. Front Comput Neurosci, 7, 163.

13. Bridge, H., Leopold, D. A. & Bourne, J. A. 2016. Adaptive Pulvinar Circuitry Supports Visual Cognition. Trends Cogn Sci, 20, 146–157.

14. Brown, E. C., Clark, D. L., Hassel, S., Macqueen, G. & Ramasubbu, R. 2017. Thalamocortical connectivity in major depressive disorder. J Affect Disord, 217, 125–131.

15. Casagrande, V. A., Sary, G., Royal, D. & Ruiz, O. 2005. On the impact of attention and motor planning on the lateral geniculate nucleus. Prog Brain Res, 149, 11–29.

16. Chai, X. J., Castanon, A. N., Ongur, D. & Whitfield-Gabrieli, S. 2012. Anticorrelations in resting state networks without global signal regression. Neuroimage, 59, 1420–8.

17. Chang, L. J., Yarkoni, T., Khaw, M. W. & Sanfey, A. G. 2013. Decoding the role of the insula in human cognition: functional parcellation and large-scale reverse inference. Cereb Cortex, 23, 739–49.

18. Charyasz, E., Heule, R., Molla, F., Erb, M., Kumar, V. J., Grodd, W., Scheffler, K. & Bause, J. 2023. Functional mapping of sensorimotor activation in the human thalamus at 9.4 Tesla. Front Neurosci, 17, 1116002.

19. Cortes, N., Ladret, H. J., Abbas-Farishta, R. & Casanova, C. 2024. The pulvinar as a hub of visual processing and cortical integration. Trends Neurosci, 47, 120–134.

20. Dacre, J., Colligan, M., Clarke, T., Ammer, J. J., Schiemann, J., Chamosa-Pino, V., Claudi, F., Harston, J. A., Eleftheriou, C., Pakan, J. M. P., Huang, C. C., Hantman, A. W., Rochefort, N. L. & Duguid, I. 2021. A cerebellar-thalamocortical pathway drives behavioral context-dependent movement initiation. Neuron, 109, 2326–2338 e8.

21. de Zeeuw, C. I. & Ten Brinke, M. M. 2015. Motor Learning and the Cerebellum. Cold Spring Harb Perspect Biol, 7, a021683.

22. Draganski, B., Kherif, F., Kloppel, S., Cook, P. A., Alexander, D. C., Parker, G. J., Deichmann, R., Ashburner, J. & Frackowiak, R. S. 2008. Evidence for segregated and integrative connectivity patterns in the human Basal Ganglia. J Neurosci, 28, 7143–52.

23. Driver, J. & Noesselt, T. 2008. Multisensory interplay reveals crossmodal influences on ’sensory-specific’ brain regions, neural responses, and judgments. Neuron, 57, 11–23.

24. Fischl, B., Sereno, M. I. & Dale, A. M. 1999. Cortical surface-based analysis. II: Inflation, flattening, and a surface-based coordinate system. Neuroimage, 9, 195–207.

25. Froesel, M., Cappe, C. & Ben Hamed, S. 2021. A multisensory perspective onto primate pulvinar functions. Neurosci Biobehav Rev, 125, 231–243.

26. Gornati, S. V., Schafer, C. B., Eelkman Rooda, O. H. J., Nigg, A. L., de Zeeuw, C. I. & Hoebeek, F. E. 2018. Differentiating Cerebellar Impact on Thalamic Nuclei. Cell Rep, 23, 2690–2704.

27. Grodd, W., Kumar, V. J., Schuz, A., Lindig, T. & Scheffler, K. 2020. The anterior and medial thalamic nuclei and the human limbic system: tracing the structural connectivity using diffusion-weighted imaging. Sci Rep, 10, 10957.

28. Haber, S. & Mcfarland, N. R. 2001. The place of the thalamus in frontal cortical-basal ganglia circuits. Neuroscientist, 7, 315–24.

29. Habig, K., Kramer, H. H., Lautenschlager, G., Walter, B. & Best, C. 2023. Processing of sensory, painful and vestibular stimuli in the thalamus. Brain Struct Funct, 228, 433–447.

30. Halassa, M. M. & Kastner, S. 2017. Thalamic functions in distributed cognitive control. Nat Neurosci, 20, 1669–1679.

31. Halliday, G. M. 2009. Thalamic changes in Parkinson’s disease. Parkinsonism Relat Disord, 15 Suppl 3, S152–5.

32. Hallquist, M. N., Hwang, K. & Luna, B. 2013. The nuisance of nuisance regression: spectral misspecification in a common approach to resting-state fMRI preprocessing reintroduces noise and obscures functional connectivity. Neuroimage, 82, 208–25.

33. Hao, L., Sheng, Z., Ruijun, W., Kun, H. Z., Peng, Z. & Yu, H. 2020. Altered Granger causality connectivity within motor-related regions of patients with Parkinson’s disease: a resting-state fMRI study. Neuroradiology, 62, 63–69.

34. Homman-Ludiye, J. & Bourne, J. A. 2019. The medial pulvinar: function, origin and association with neurodevelopmental disorders. J Anat, 235, 507–520.

35. Hong, S. B. 2023. Thalamocortical functional connectivity in youth with attention-deficit/hyperactivity disorder. J Psychiatry Neurosci, 48, E50–E60.

36. Hwang, K., Bertolero, M. A., Liu, W. B. & D’esposito, M. 2017. The Human Thalamus Is an Integrative Hub for Functional Brain Networks. J Neurosci, 37, 5594–5607.

37. Iglesias, J. E., Insausti, R., Lerma-Usabiaga, G., Bocchetta, M., van Leemput, K., Greve, D. N., van der Kouwe, A., Alzheimer’S DISEASE Neuroimaging, I., Fischl, B., Caballero-Gaudes, C. & Paz-Alonso, P. M. 2018. A probabilistic atlas of the human thalamic nuclei combining ex vivo MRI and histology. Neuroimage, 183, 314–326.

38. Ilinsky, I., Horn, A., Paul-Gilloteaux, P., Gressens, P., Verney, C. & Kultas-Ilinsky, K. 2018. Human Motor Thalamus Reconstructed in 3D from Continuous Sagittal Sections with Identified Subcortical Afferent Territories. eNeuro, 5.

39. Ilyas, A., Pizarro, D., Romeo, A. K., Riley, K. O. & Pati, S. 2019. The centromedian nucleus: Anatomy, physiology, and clinical implications. J Clin Neurosci, 63, 1–7.

40. Jang, H., Plis, S. M., Calhoun, V. D. & Lee, J. H. 2017. Task-specific feature extraction and classification of fMRI volumes using a deep neural network initialized with a deep belief network: Evaluation using sensorimotor tasks. Neuroimage, 145, 314–328.

41. Jang, S. H., Lim, H. W. & Yeo, S. S. 2014. The neural connectivity of the intralaminar thalamic nuclei in the human brain: a diffusion tensor tractography study. Neurosci Lett, 579, 140–4.

42. Jankowski, M. M., Ronnqvist, K. C., Tsanov, M., Vann, S. D., Wright, N. F., Erichsen, J. T., Aggleton, J. P. & O’mara, S. M. 2013. The anterior thalamus provides a subcortical circuit supporting memory and spatial navigation. Front Syst Neurosci, 7, 45.

43. Jarrett, C., Zwosta, K., Wang, X., Wolfensteller, U., Iglesias, J. E., VON Kriegstein, K. & Ruge, H. 2024. Progressive Changes in Functional Connectivity between Thalamic Nuclei and Cortical Networks Across Learning. bioRxiv, 2024.08.26.609333.

44. Jones, E. G. 1983. The Thalamus, New York, Springer Science+Business Media, LLC.

45. Jones, E. G. 1998. A new view of specific and nonspecific thalamocortical connections. Adv Neurol, 77, 49–71; discussion 72-3.

46. Jones, E. G. 2007. The Thalamus 2 Volume Set, Cambridge University Press.

47. Kark, S. M., Birnie, M. T., Baram, T. Z. & Yassa, M. A. 2021. Functional Connectivity of the Human Paraventricular Thalamic Nucleus: Insights From High Field Functional MRI. Front Integr Neurosci, 15, 662293.

48. Kerestes, C., Delafield, R., Elia, J., Shochet, T., Kaneshiro, B. & Soon, R. 2022. Person-centered, high-quality care from a distance: A qualitative study of patient experiences of TelAbortion, a model for direct-to-patient medication abortion by mail in the United States. Perspect Sex Reprod Health, 54, 177–187.

49. Koster, K. P. & Sherman, S. M. 2024. Convergence of inputs from the basal ganglia with layer 5 of motor cortex and cerebellum in mouse motor thalamus. Elife, 13.

50. Kowalczyk, O. S., Mehta, M. A., O’daly, O. G. & Criaud, M. 2022. Task-Based Functional Connectivity in Attention-Deficit/Hyperactivity Disorder: A Systematic Review. Biol Psychiatry Glob Open Sci, 2, 350–367.

51. Kumar, V. J., Beckmann, C. F., Scheffler, K. & Grodd, W. 2022. Relay and higher-order thalamic nuclei show an intertwined functional association with cortical-networks. Commun Biol, 5, 1187.

52. Kumar, V. J., Scheffler, K. & Grodd, W. 2023. The structural connectivity mapping of the intralaminar thalamic nuclei. Sci Rep, 13, 11938.

53. Kumar, V. J., Van Oort, E., Scheffler, K., Beckmann, C. F. & Grodd, W. 2017. Functional anatomy of the human thalamus at rest. Neuroimage, 147, 678–691.

54. Lambert, C., Simon, H., Colman, J. & Barrick, T. R. 2017. Defining thalamic nuclei and topographic connectivity gradients in vivo. Neuroimage, 158, 466–479.

55. Lee, S., Kim, J. & Tak, S. 2020. Effects of Autonomous Sensory Meridian Response on the Functional Connectivity as Measured by Functional Magnetic Resonance Imaging. Front Behav Neurosci, 14, 154.

56. Lesica, N. A. & Stanley, G. B. An LGN Inspired Detect/Transmit Framework for High Fidelity Relay of Visual Information with Limited Bandwidth. 2005 Berlin, Heidelberg. Springer Berlin Heidelberg, 177–186.

57. Manto, M., Bower, J. M., Conforto, A. B., Delgado-Garcia, J. M., da Guarda, S. N., Gerwig, M., Habas, C., Hagura, N., Ivry, R. B., Marien, P., Molinari, M., Naito, E., Nowak, D. A., Oulad Ben Taib, N., Pelisson, D., Tesche, C. D., Tilikete, C. & Timmann, D. 2012. Consensus paper: roles of the cerebellum in motor control--the diversity of ideas on cerebellar involvement in movement. Cerebellum, 11, 457–87.

58. Mastropasqua, C., Bozzali, M., Spano, B., Koch, G. & Cercignani, M. 2015. Functional Anatomy of the Thalamus as a Model of Integrated Structural and Functional Connectivity of the Human Brain In Vivo. Brain Topogr, 28, 548–58.

59. Mcfarland, N. R. & Haber, S. N. 2002. Thalamic relay nuclei of the basal ganglia form both reciprocal and nonreciprocal cortical connections, linking multiple frontal cortical areas. J Neurosci, 22, 8117–32.

60. Mehler, D. M. A. & Reschechtko, S. 2018. Movement Variability Is Processed Bilaterally by Inferior Parietal Lobule. J Neurosci, 38, 2413–2415.

61. Meng, Q. & Schneider, K. A. 2022. The magnocellular division of the human medial geniculate nucleus preferentially responds to auditory transients. bioRxiv, 2022.06.05.494907.

62. Middleton, F. A. & Strick, P. L. 2000. Basal ganglia and cerebellar loops: motor and cognitive circuits. Brain Res Brain Res Rev, 31, 236–50.

63. Mitchell, A. S. 2015. The mediodorsal thalamus as a higher order thalamic relay nucleus important for learning and decision-making. Neurosci Biobehav Rev, 54, 76–88.

64. Moeller, S., Pisharady, P. K., Ramanna, S., Lenglet, C., Wu, X., Dowdle, L., Yacoub, E., Ugurbil, K. & Akcakaya, M. 2021. NOise reduction with DIstribution Corrected (NORDIC) PCA in dMRI with complex-valued parameter-free locally low-rank processing. Neuroimage, 226, 117539.

65. Mohagheghi Nejad, M., Rotter, S. & Schmidt, R. 2018. Transmission of motor signals from the basal ganglia to the thalamus: effect of correlations, sensory responses, and excitation. bioRxiv, 386920.

66. Nagalski, A., Puelles, L., Dabrowski, M., Wegierski, T., Kuznicki, J. & Wisniewska, M. B. 2016. Molecular anatomy of the thalamic complex and the underlying transcription factors. Brain Struct Funct, 221, 2493–510.

67. Nelson, A. J. D. 2021. The anterior thalamic nuclei and cognition: A role beyond space? Neurosci Biobehav Rev, 126, 1–11.

68. Nieto-Castanon, A. 2020. FMRI minimal preprocessing pipeline. Handbook of functional connectivity Magnetic Resonance Imaging methods in CONN. Hilbert Press.

69. Nieto-Castanon, A. 2020b. Functional connectivity measures. Handbook of functional connectivity magnetic resonance imaging methods in CONN. Hilbert Press.

70. Nieto-Castanon, A. & Whitfield-Gabrieli, S. 2022. CONN functional connectivity toolbox (RRID: SCR_009550), Version 22, Hilbert Press.

71. O’connor, K. N., Allison, T. L., Rosenfield, M. E. & Moore, J. W. 1997. Neural activity in the medial geniculate nucleus during auditory trace conditioning. Exp Brain Res, 113, 534–56.

72. Owens-Walton, C., Jakabek, D., Power, B. D., Walterfang, M., Velakoulis, D., Van Westen, D., Looi, J. C. L., Shaw, M. & Hansson, O. 2019. Increased functional connectivity of thalamic subdivisions in patients with Parkinson’s disease. PLoS One, 14, e0222002.

73. Palejwala, A. H., Dadario, N. B., Young, I. M., O’connor, K., Briggs, R. G., Conner, A. K., O’donoghue, D. L. & Sughrue, M. E. 2021. Anatomy and White Matter Connections of the Lingual Gyrus and Cuneus. World Neurosurg, 151, e426–e437.

74. Palesi, F., Tournier, J. D., Calamante, F., Muhlert, N., Castellazzi, G., Chard, D., D’angelo, E. & Wheeler-Kingshott, C. A. 2015. Contralateral cerebello-thalamo-cortical pathways with prominent involvement of associative areas in humans in vivo. Brain Struct Funct, 220, 3369–84.

75. Papez, J. W. 1995. A proposed mechanism of emotion. 1937. J Neuropsychiatry Clin Neurosci, 7, 103–12.

76. Pisano, T. J., Dhanerawala, Z. M., Kislin, M., Bakshinskaya, D., Engel, E. A., Hansen, E. J., Hoag, A. T., Lee, J., de Oude, N. L., Venkataraju, K. U., Verpeut, J. L., Hoebeek, F. E., Richardson, B. D., Boele, H. J. & Wang, S. S. 2021. Homologous organization of cerebellar pathways to sensory, motor, and associative forebrain. Cell Rep, 36, 109721.

77. Popa, L. S., Streng, M. L., Hewitt, A. L. & Ebner, T. J. 2016. The Errors of Our Ways: Understanding Error Representations in Cerebellar-Dependent Motor Learning. Cerebellum, 15, 93–103.

78. Prati, J. M., Pontes-Silva, A. & Gianlorenco, A. C. L. 2024. The cerebellum and its connections to other brain structures involved in motor and non-motor functions: A comprehensive review. Behav Brain Res, 465, 114933.

79. Rodriguez-Sabate, C., Llanos, C., Morales, I., Garcia-Alvarez, R., Sabate, M. & Rodriguez, M. 2015. The functional connectivity of intralaminar thalamic nuclei in the human basal ganglia. Hum Brain Mapp, 36, 1335–47.

80. Rolls, E. T. 2016. Cerebral Cortex: Principles of Operation, Oxford University Press.

81. Rudolph, S., Badura, A., Lutzu, S., Pathak, S. S., Thieme, A., Verpeut, J. L., Wagner, M. J., Yang, Y.-M. & Fioravante, D. 2023. Cognitive-Affective Functions of the Cerebellum. The Journal of Neuroscience, 43, 7554–7564.

82. Saalmann, Y. B. 2014. Intralaminar and medial thalamic influence on cortical synchrony, information transmission and cognition. Front Syst Neurosci, 8, 83.

83. Saalmann, Y. B. & Kastner, S. 2011. Cognitive and perceptual functions of the visual thalamus. Neuron, 71, 209–23.

84. Saalmann, Y. B. & Kastner, S. 2015. The cognitive thalamus. Frontiers in Systems Neuroscience, 9.

85. Saalmann, Y. B., Pinsk, M. A., Wang, L., Li, X. & Kastner, S. 2012. The pulvinar regulates information transmission between cortical areas based on attention demands. Science, 337, 753–6.

86. Sasaki, R., Kumano, H., Mitani, A., Suda, Y. & Uka, T. 2022. Task-specific employment of sensory signals underlies rapid task switching. Cereb Cortex, 32, 4657–4670.

87. Semrau, J. A., Herter, T. M., Kiss, Z. H. & Dukelow, S. P. 2015. Disruption in proprioception from long-term thalamic deep brain stimulation: a pilot study. Front Hum Neurosci, 9, 244.

88. Shajan, G., Kozlov, M., Hoffmann, J., Turner, R., Scheffler, K. & Pohmann, R. 2014. A 16-channel dual-row transmit array in combination with a 31-element receive array for human brain imaging at 9.4 T. Magn Reson Med, 71, 870–9.

89. Sherman, S. M. 2007. The thalamus is more than just a relay. Curr Opin Neurobiol, 17, 417–22.

90. Sherman, S. M. 2016. Thalamus plays a central role in ongoing cortical functioning. Nat Neurosci, 19, 533–41.

91. Sherman, S. M. & Guillery, R. W. 2006. Exploring the thalamus and its role in cortical function, MIT press.

92. Shimono, M., Mano, H. & Niki, K. 2012. The brain structural hub of interhemispheric information integration for visual motion perception. Cereb Cortex, 22, 337–44.

93. Shine, J. M., Lewis, L. D., Garrett, D. D. & Hwang, K. 2023. The impact of the human thalamus on brain-wide information processing. Nat Rev Neurosci, 24, 416–430.

94. Smith, S. M., Jenkinson, M., Woolrich, M. W., Beckmann, C. F., Behrens, T. E., Johansen-Berg, H., Bannister, P. R., de Luca, M., Drobnjak, I., Flitney, D. E., Niazy, R. K., Saunders, J., Vickers, J., Zhang, Y., de Stefano, N., Brady, J. M. & Matthews, P. M. 2004. Advances in functional and structural MR image analysis and implementation as FSL. Neuroimage, 23 Suppl 1, S208–19.

95. Smith, Y., Raju, D., Nanda, B., Pare, J. F., Galvan, A. & Wichmann, T. 2009. The thalamostriatal systems: anatomical and functional organization in normal and parkinsonian states. Brain Res Bull, 78, 60–8.

96. Sommer, M. A. 2003. The role of the thalamus in motor control. Curr Opin Neurobiol, 13, 663–70.

97. Spets, D. S. & Slotnick, S. D. 2020. Thalamic Functional Connectivity during Spatial Long-Term Memory and the Role of Sex. Brain Sci, 10.

98. Stoodley, C. J., Valera, E. M. & Schmahmann, J. D. 2012. Functional topography of the cerebellum for motor and cognitive tasks: an fMRI study. Neuroimage, 59, 1560–70.

99. Studtmann, C., Ladislav, M., Safari, M., Khondaker, R., Chen, Y., Vaughan, G. A., Topolski, M. A., Tomovic, E., Balik, A. & Swanger, S. A. 2023. Ventral posterolateral and ventral posteromedial thalamocortical neurons have distinct physiological properties. J Neurophysiol, 130, 1492–1507.

100. Sun, N., Liu, M., Liu, P., Zhang, A., Yang, C., Liu, Z., Li, J., Li, G., Wang, Y. & Zhang, K. 2023. Abnormal cortical-striatal-thalamic-cortical circuit centered on the thalamus in MDD patients with somatic symptoms: Evidence from the REST-meta-MDD project. J Affect Disord, 323, 71–84.

101. Sweeney-Reed, C. M., Zaehle, T., Voges, J., Schmitt, F. C., Buentjen, L., Borchardt, V., Walter, M., Hinrichs, H., Heinze, H. J., Rugg, M. D. & Knight, R. T. 2017. Anterior Thalamic High Frequency Band Activity Is Coupled with Theta Oscillations at Rest. Front Hum Neurosci, 11, 358.

102. Tomasi, D., Wang, R., Wang, G. J. & Volkow, N. D. 2014. Functional connectivity and brain activation: a synergistic approach. Cereb Cortex, 24, 2619–29.

103. van der Werf, Y. D., Witter, M. P. & Groenewegen, H. J. 2002. The intralaminar and midline nuclei of the thalamus. Anatomical and functional evidence for participation in processes of arousal and awareness. Brain Res Brain Res Rev, 39, 107–40.

104. Wagner, G., de la Cruz, F., Schachtzabel, C., Gullmar, D., Schultz, C. C., Schlosser, R. G., Bar, K. J. & Koch, K. 2015. Structural and functional dysconnectivity of the fronto-thalamic system in schizophrenia: a DCM-DTI study. Cortex, 66, 35–45.

105. Wahlbom, A., Enander, J. M. D. & Jorntell, H. 2021. Widespread Decoding of Tactile Input Patterns Among Thalamic Neurons. Front Syst Neurosci, 15, 640085.

106. Wang, S., Cai, H., Cao, Z., Li, C., Wu, T., Xu, F., Qian, Y., Chen, X. & Yu, Y. 2021. More Than Just Static: Dynamic Functional Connectivity Changes of the Thalamic Nuclei to Cortex in Parkinson’s Disease With Freezing of Gait. Frontiers in Neurology, 12.

107. Wang, Y. 2024. Thalamus and its functional connections with cortical regions contribute to complexity-dependent cognitive reasoning. Neuroscience, 562, 125–134.

108. Ward, L. M. 2013. The thalamus: gateway to the mind. Wiley Interdiscip Rev Cogn Sci, 4, 609–622.

109. Whitfield-Gabrieli, S. & Nieto-Castanon, A. 2012. Conn: a functional connectivity toolbox for correlated and anticorrelated brain networks. Brain Connect, 2, 125–41.

110. Wilke, M., Schneider, L., Dominguez-Vargas, A. U., Schmidt-Samoa, C., Miloserdov, K., Nazzal, A., Dechent, P., Cabral-Calderin, Y., Scherberger, H., Kagan, I. & Bahr, M. 2018. Reach and grasp deficits following damage to the dorsal pulvinar. Cortex, 99, 135–149.

111. Woodward, N. D., Karbasforoushan, H. & Heckers, S. 2012. Thalamocortical dysconnectivity in schizophrenia. Am J Psychiatry, 169, 1092–9.

112. Wright, N. F., Vann, S. D., Aggleton, J. P. & Nelson, A. J. 2015. A critical role for the anterior thalamus in directing attention to task-relevant stimuli. J Neurosci, 35, 5480–8.

113. Zhang, D., Snyder, A. Z., Fox, M. D., Sansbury, M. W., Shimony, J. S. & Raichle, M. E. 2008. Intrinsic functional relations between human cerebral cortex and thalamus. J Neurophysiol, 100, 1740–8.

114. Zhang, D., Snyder, A. Z., Shimony, J. S., Fox, M. D. & Raichle, M. E. 2010. Noninvasive functional and structural connectivity mapping of the human thalamocortical system. Cereb Cortex, 20, 1187–94.

115. Zhang, J., Chu, K. W., Teague, E. B., Newmark, R. E. & Buchsbaum, M. S. 2013. fMRI assessment of thalamocortical connectivity during attentional performance. Magn Reson Imaging, 31, 1112–8.

116. Zhou, H., Schafer, R. J. & Desimone, R. 2016. Pulvinar-Cortex Interactions in Vision and Attention. Neuron, 89, 209–20.

117. Zhou, J., Liu, X., Song, W., Yang, Y., Zhao, Z., Ling, F., Hudetz, A. G. & Li, S. J. 2011. Specific and nonspecific thalamocortical functional connectivity in normal and vegetative states. Conscious Cogn, 20, 257–68.

