## Supplementary material for "Functional connectivity of thalamic nuclei during sensorimotor task-based fMRI at 9.4 Tesla": Suplemental Figures

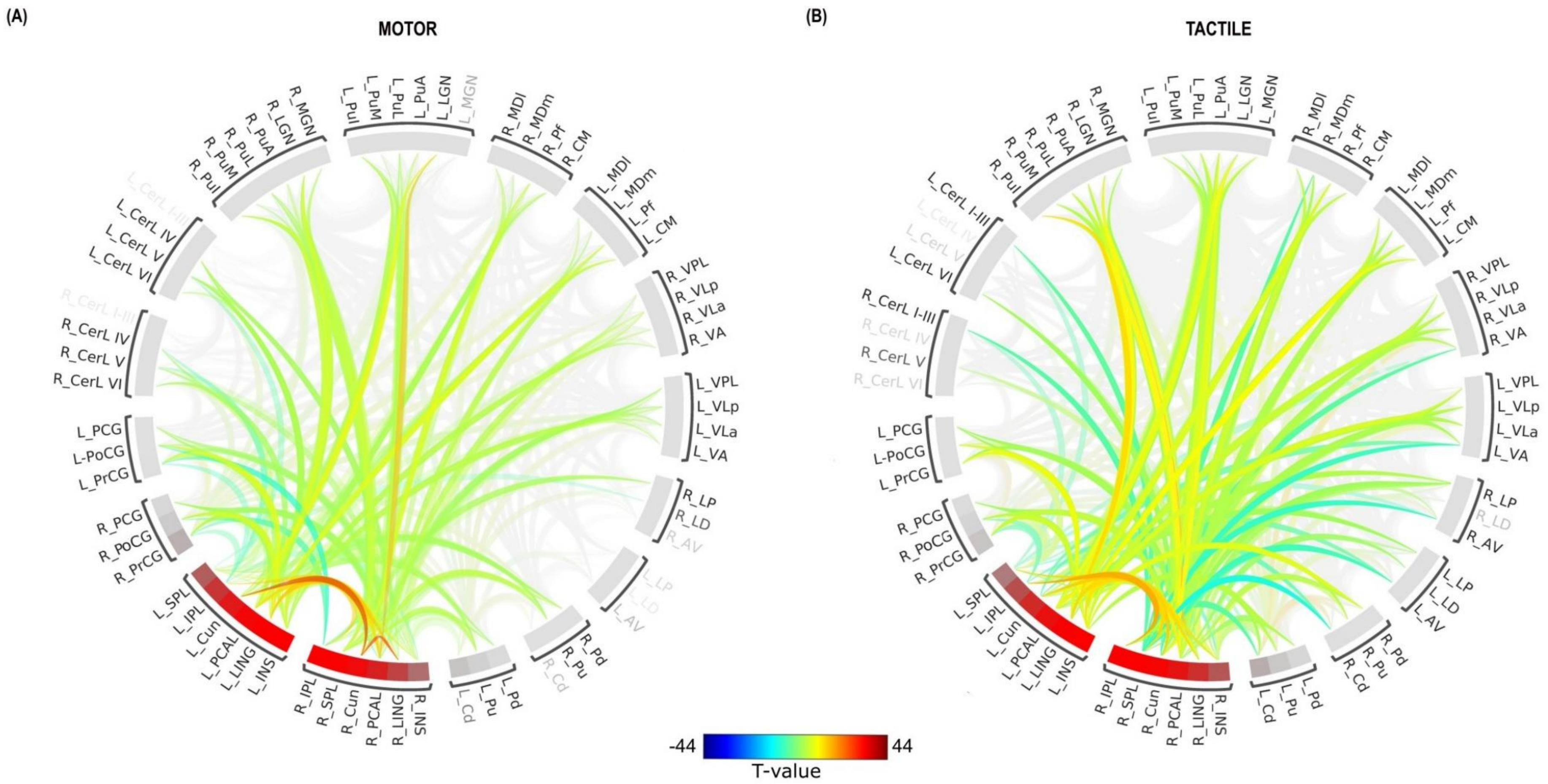


Figure S1. ROI-to-ROI connectome ring of functional connectivity between cortical and other brain regions for the motor (A) and tactile (B) tasks. The color links represent ROI-to-ROI connections with a false discovery rate (FDR) of $p < 0.05$. Warm colors (yellow and red) indicate increased connectivity, while cool colors (dark and light blue) signify decreased connectivity.

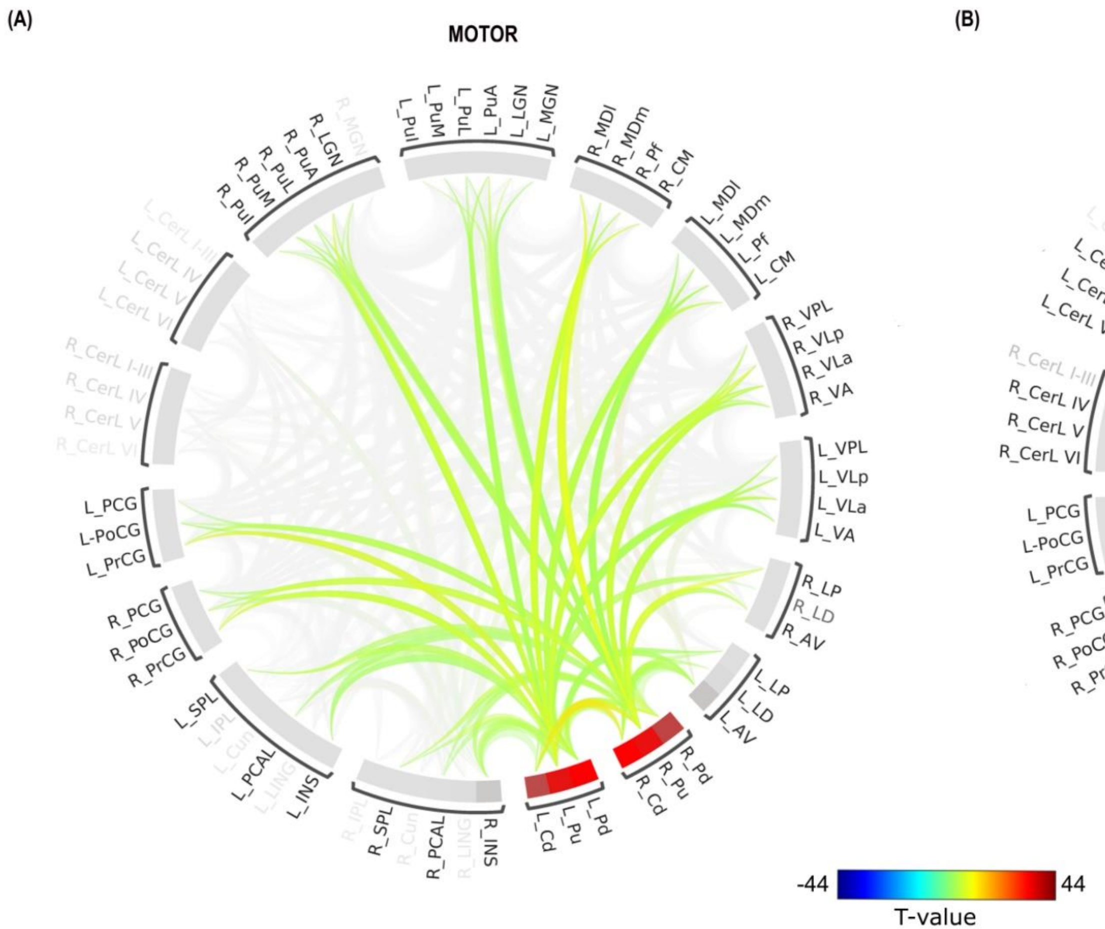


Figure S2. ROI-to-ROI connectome ring of functional connectivity between basal ganglia nuclei and other brain regions for the motor (A) and tactile (B) tasks. The color links represent ROI-to-ROI connections with a false discovery rate (FDR) of $p < 0.05$. Warm colors (yellow and red) indicate increased connectivity, while cool colors (dark and light blue) signify decreased connectivity.

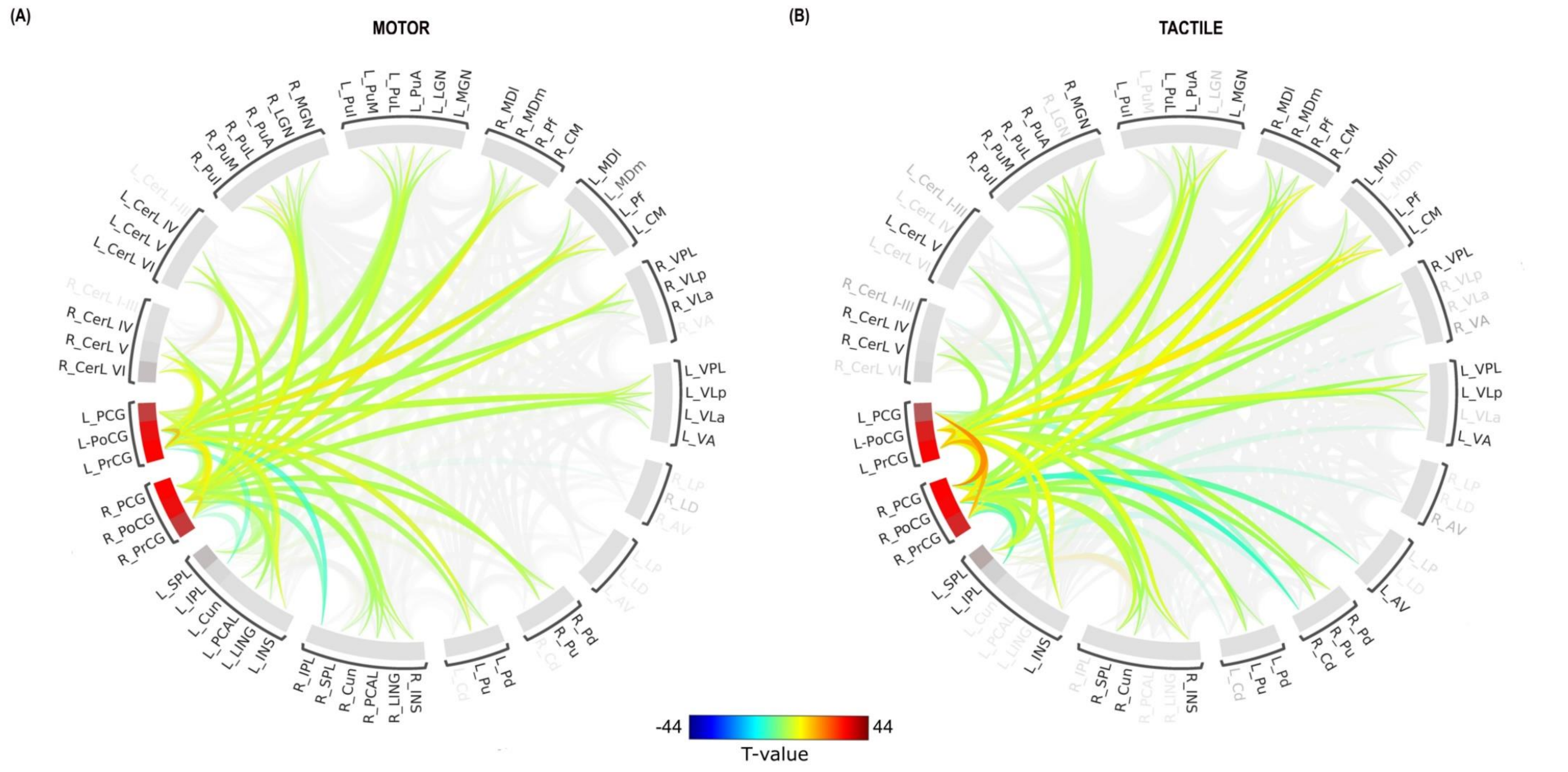


Figure S3. ROI-to-ROI connectome ring of functional connectivity between sensorimotor cortical regions and other brain regions for the motor (A) and tactile (B) tasks. The color links represent ROI-to-ROI connections with a false discovery rate (FDR) of $p < 0.05$. Warm colors (yellow and red) indicate increased connectivity, while cool colors (dark and light blue) signify decreased connectivity.

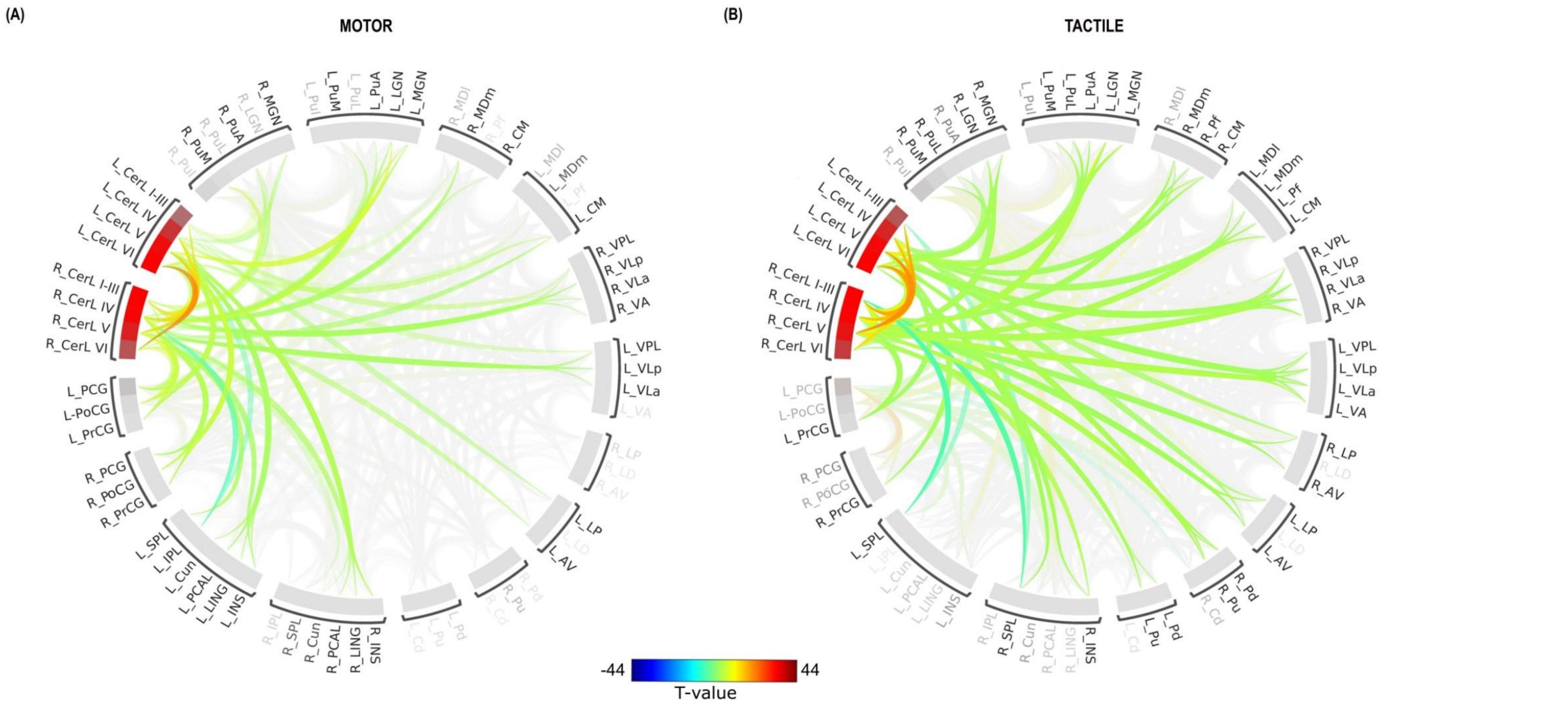


Figure S4. ROI-to-ROI connectome ring of functional connectivity between cerebellar regions and other brain regions for the motor (A) and tactile (B) tasks. The color links represent ROI-to-ROI connections with a false discovery rate (FDR) of $p < 0.05$. Warm colors (yellow and red) indicate increased connectivity, while cool colors (dark and light blue) signify decreased connectivity.
